# Default template search through the ColabFold server supplies OpenFold3 benchmark targets with their own structures: frequency and bounds on the accuracy effect

**DOI:** 10.64898/2026.11.24.769011

**Authors:** Mahdi Soultana, Olamide Isreal Odesanya, Mervin Gaitho, Gabriel Richman

## Abstract

Protein structure prediction models are tested on structures released after their training data, but at prediction time they also search a database of known structures for templates, and that database can contain the test structures themselves. We measured how often this happens for OpenFold3 0.4.2 run with its default template search through the ColabFold MMseqs2 server, and whether it changes accuracy. Of 61 X-ray structures released after the 2021-09-30 training cutoff, only the 30 released on or before 17 May 2023, the inferred date of the server’s template database, could retrieve their own entry. For 16 of these 30, the default search supplied the target’s own entry or an entry released the same day with at least 99.3% sequence identity. We predicted all 61 targets (monomers and protein, small-molecule and nucleic-acid complexes) with the default search, with templates limited to the cutoff, without templates, and with only own and same-day entries removed; all settings of a category ran in one job, on one graphics card, from one set of alignments and template hits. The accuracy comparison bounds an effect rather than measuring it. Limiting templates to the cutoff changed monomer accuracy by a median of 0.036 Å (adjusted p = 0.11), and with 15 monomers only shifts of about 0.26 Å were detectable; without the seven targets used during pipeline development the median was 0.041 Å (unadjusted p = 0.005). Interface accuracy of complexes was equivalent within 0.05 DockQ or 0.5 Å; templates act per protein chain, so partner placement was not tested. Under this configuration, exposure of benchmark targets to their own structures is common, and its average effect was below what this target set can detect. Date-based benchmarks should limit templates to the training cutoff and report which templates were supplied.

## Introduction

AlphaFold 3 [1] and open reproductions such as OpenFold3 [2], Boltz-1 [3] and Protenix [4] predict complexes of proteins, nucleic acids and small molecules. Their accuracy is usually measured on PDB entries released after the training cutoff. The AlphaFold 3 evaluation limited templates to entries released before the training cutoff [1]; OpenFold3 as run here does not apply this limit by default (Materials and methods), and a later release date does not imply that a target is unrelated to earlier structures. Runs N’ Poses, a set of 2,600 protein–ligand systems released after the training cutoff, showed that four co-folding methods largely reproduce ligand poses seen in training [5, 14], and regularly updated benchmark sets have been proposed to limit leakage in AlphaFold benchmarks more generally [6]. Leakage-aware datasets such as PLINDER [7] split by structural and sequence similarity rather than by date.

Three routes can carry information about a post-cutoff target into a prediction: related structures in the training data, templates supplied at inference, and multiple sequence alignments built from databases that postdate the cutoff. This study measures the effect of the second route on OpenFold3 across four complex types, reports sequence identity to the pre-cutoff PDB as a proxy for the first, and does not measure the third.

Varying AlphaFold 3 template and MSA inputs changes accuracy on CASP16 targets [8], and FoldBench evaluates all-atom predictors on low-homology assemblies [9]. Those studies restrict or perturb the inputs. Limiting templates by release date is established practice: the AlphaFold2 pipeline takes a maximum template release date as a run setting [17], and the AlphaFold 3 evaluation applied one [1]. Runs N’ Poses ran AlphaFold 3 with and without templates and found no significant difference in ligand accuracy, which its authors attribute to template ligand information not being used [14]. None of [8], [9] and [14] reports how often default inference retrieves the target’s own entry and supplies it to the model, and no such report was found in Europe PMC and web searches (2026-11-20) on template leakage and own-entry templates for AlphaFold 3, Boltz, Chai and OpenFold3. This study asks how often default OpenFold3 inference retrieves and supplies post-cutoff or own-entry structures for a date-based test set, and whether that exposure changes accuracy. The template database searched at inference is set by the inference service rather than fixed by the model.

Default OpenFold3 inference supplied the target’s own entry or a same-day entry to the model as a template for 16 of the 30 targets that its inferred template database could contain, and retrieved one for 21. We report this exposure by complex type, at the level of the template chains passed to the model, the accuracy of the same targets under unrestricted, cutoff-limited and no templates, their sequence relatedness to pre-cutoff structures, and the variation between seeds.

## Materials and methods

### Target set

Candidates were retrieved from the RCSB PDB with four queries for X-ray entries at 2.5 Å resolution or better released from 2023-01-01 to 2024-06-30. Each query returned its first 400 hits by release date (800 for ligands), so the sampling frame is weighted toward 2023: 53 of the 61 targets were released in 2023 and 30 on or before 2023-05-17 (Table S1).

- **Monomers:** one protein entity, one instance, 80 to 400 residues.
- **Protein–protein:** two protein entities, one instance each, 150 to 600 residues in total, each chain at least 40 residues.
- **Protein–ligand:** one protein entity of 120 to 400 residues with exactly one non-polymer component of 250 to 700 Da, present once, excluding common additives and cofactors and entries titled as covalent complexes.
- **Protein–nucleic acid:** one protein entity of at least 60 residues with one or two DNA or RNA entities of at least 6 nucleotides, 60 to 450 residues in total, at most three instances.

Candidates were shuffled with seed 20261118 and accepted unless a protein chain shared 30% or more sequence identity (local alignment, BLOSUM62, identity over the shorter sequence) with an accepted chain in any category, or the entry contained non-standard residues. 8GAR (covalently bound heme) and 8AAE (carborane ligand) were excluded. The selection as made is archived (original_selection/, with the seven targets also used for pipeline development: 8T7K, 8T2D, 8IF0, 8BM8, 8RU8, 7WKP, 8B4C), and verify_cohort.py checks every target against it. Re-running the selection now picks a different sample, because the program shuffles the first 400 hits by release date and the content of that window has changed; the original candidate lists were not retained (Supplementary text S1).

### Relatedness to the pre-cutoff PDB

For each protein chain, the highest sequence identity to a PDB polymer entity released on or before 2021-09-30 was taken from the RCSB sequence search service (MMseqs2, E-value 1). Identity to the pre-cutoff PDB indicates relatedness to structures that were available for training; it does not establish training-set membership.

For each own-entry or same-day template chain, the chain sequence (by label_asym_id, the chain naming OpenFold3 uses for templates) was taken from the RCSB Data API, and identity and coverage of the target chain it was retrieved for were computed with the local alignment used for target selection (BLOSUM62, gap opening −11, extension −1, identity over the shorter sequence).

### Predictions

OpenFold3 0.4.2 with the of3-p2-155k checkpoint was run in a Nextflow pipeline using container harbor.cluster.omic.ai/omic/openfold_3@sha256:8417e338eeb0931224455b743a367e8a5a f4003b76d6bd877bee4832732a2a74, on an internal registry; the pipeline and a build recipe that uses only public inputs (OpenFold3 source with its dependency lock file, and the checkpoint from the public AWS Open Data bucket, checked by sha256) are deposited (openfold3_pipeline/). The OpenFold3 technical report evaluates held-out performance against the AlphaFold 3 training cutoff of 2021-09-30 [2], and this date is referred to as the training cutoff throughout. Inputs were canonical SEQRES sequences and, for ligands, stereo SMILES from the PDB Chemical Component Dictionary. MSAs were built with the ColabFold MMseqs2 server. DeepSpeed evoformer attention and cuEquivariance kernels were disabled.

Each category was predicted in one Nextflow task (openfold_3_summary, TRS version change-3), so all template settings of a category ran on one GPU, one container and one pipeline commit. The task first ran OpenFold3 with the ColabFold MMseqs2 server for one seed, which fetched the MSAs and the raw template hit list of every protein chain; these server responses are archived (v16_msa_cache/ in the data deposit [16]). Each template setting was then run with three seeds and five diffusion samples from that saved query set, with the MSA server off, so all settings read the same MSAs and the same hit lists. OpenFold3 template preprocessing ran again for each setting in its own cache directory, because OpenFold3 0.4.2 reuses a cached template list without re-applying the release-date filter. Template settings were:

1. **Unrestricted:** default template search.
2. **Cutoff:** templates released on or after 2021-09-30 removed by OpenFold3 (template_preprocessor_settings.max_release_date).
3. **None:** use_templates = false.
4. **Self-excluded:** the hit list of each protein chain without the target’s own entry and without entries released on the target’s release date; every other hit kept its position.
5. **Cutoff without MSA (monomers only):** setting 2 with no MSA, to compare the template effect with the contribution of the MSA.

The setting that reused the saved inputs reproduced the server setting’s template list for 76 of 77 protein chains (for 8GTI chain A the server setting kept no template and every reused setting kept 20 from the same hit list; the server run’s template preprocessing for that chain did not complete, and all reported 8GTI contrasts use the reused settings, which are consistent with one another) and its model within 0.007 Å Cα RMSD for the same seed in a pilot target. Monomers ran on one RTX 3090 and the three complex categories on a second RTX 3090 (gpu.txt). The ligand centroid distances used a separate set of single-seed predictions (Table S1).

No template retrieved under the cutoff setting had a recorded release date after 2021-09-30. Three retrieved templates (1JT1, 6TTL, 6WBO) are obsolete entries without a recorded release date. By default, OpenFold3 obtains template hits from the ColabFold MSA server [10], whose template database is listed as PDB100 230517; MSAs from the same server use UniRef30 2302 and ColabFoldDB 202108 [11]. The latest release date observed among unrestricted templates, 2023-05-17, matches this database version. If the server used PDB100 230517, the 31 targets released after that date could not have retrieved their own entry. For each protein chain, OpenFold3 0.4.2 records in its processed query set the template chains that passed template preprocessing, in the order of the server’s hit list, and at inference supplies the first four of them to the model (default n_templates = 4 with top-k selection). A template is called retrieved when it is on that list and supplied when it is among the first four. Templates under the unrestricted setting were classed as post-cutoff (release date after 2021-09-30) and, more strictly, as self-related (the target’s own identifier or an entry released on the same day as the target).

### Software versions

OpenFold3 0.4.2 ran with torch 2.10.0 and cuEquivariance 0.10.0 under Nextflow 26.10.2; scoring used biopython 1.88, DockQ 2.1.3, PoseBusters 0.6.5, RDKit 2026.9.2, scipy 1.17.1 and statsmodels 0.15.0. The full list is in Supplementary text S3 and software_versions.json.

### Scoring

For each seed, the sample ranked first by OpenFold3’s sample ranking score was compared with the first model of the deposited structure using label sequence numbering, and per-target values are the mean over three seeds (scoring provenance in Supplementary text S3).

- **Monomers:** Cα RMSD after global superposition; TM-score normalized by reference length.
- **Interfaces:** DockQ v2 [12] (DockQ 2.1.3, Python API) of the top-ranked model against the deposited structure, with the identity chain mapping (model chains carry the reference chain names) and every native interface between polymer chains scored; DockQ combines the fraction of native contacts, interface RMSD and ligand RMSD. The protein–protein value is the DockQ of the protein–protein interface; the protein–nucleic-acid value is the mean DockQ over the protein–nucleic-acid interfaces with native contacts, excluding nucleic-acid–nucleic-acid interfaces. DockQ thresholds (0.23 acceptable, 0.49 medium) were defined for protein–protein interfaces and are applied unchanged to protein–nucleic-acid interfaces.
- **Protein–ligand:** pocket Cα atoms (residues within 10 Å of the reference ligand) were superposed, and the symmetry-corrected heavy-atom ligand RMSD was computed with RDKit after assigning bond orders from the input SMILES. Ligand centroid displacement after global superposition was computed to identify wrong-site predictions. Pose validity was assessed with PoseBusters [13] in redock mode against the predicted receptor. A pose was counted as valid when it passed all redock checks except the reference-pose RMSD check, which measures pose accuracy rather than chemical or geometric validity.

Success thresholds were Cα RMSD below 2 Å, interface DockQ of at least 0.23, and ligand RMSD below 2 Å.

### Statistics

The target was the unit of inference, and each target’s value was the mean over three seeds of the top-ranked prediction. The primary family was the three setting pairs (unrestricted vs cutoff, cutoff vs none, unrestricted vs none) in four categories; two-sided exact Wilcoxon signed-rank tests (zero differences dropped, all pairs were nonzero) were Holm-adjusted over these 12 tests. Median paired differences are given with distribution-free order-statistic intervals of at least 95% coverage. Equivalence was tested with two one-sided Wilcoxon tests (TOST) at margins of 0.1 Å Cα RMSD, 0.05 DockQ and 0.5 Å ligand RMSD. The margins were fixed before the reported runs but after earlier runs of the same targets had been analysed (Supporting information); for Cα RMSD and DockQ they are about three times the largest median seed-to-seed standard deviation observed in those runs (0.036 Å and 0.017 DockQ; Table S8), the ligand margin is a quarter of the 2 Å success threshold, and none was justified by a formal criterion. The study was not preregistered; the self-excluded setting, the template TM-scores and the MSA control were added after the earlier runs were reviewed, and are reported as exploratory. For each contrast the smallest median shift detected with 80% power was estimated by shifting the centred observed differences and resampling their signs (2,000 draws). The self-exposed subsets and the self-excluded setting form a second, exploratory family, Holm-adjusted within itself, and the MSA contrast a third. Proportions are given with Wilson 95% confidence intervals. As a noise floor, the unrestricted and self-excluded settings receive identical template lists for targets with no own or same-day hit, so their difference measures run-to-run variation on these inputs. Results without the seven development targets are given for every category. All numbers are produced by analysis_v16.py, extra_v16.py and tables_v16.py from the score files. One protein–ligand prediction (8CJI, one seed, unrestricted setting) could not be scored, so 8CJI is excluded from the contrasts that involve the unrestricted setting (n = 13) and kept in the cutoff-versus-none contrast (n = 14).

### Use of AI tools

The study was run on the Omic research platform (https://omic.ai). AI agents on the platform, built on large language models (Claude, Anthropic, including Claude Opus 5.5 through Claude Code; and Codex, OpenAI), wrote the target-selection, prediction-pipeline, scoring and statistics scripts, ran the predictions and analyses, and drafted the manuscript text under the authors’ direction. Claude Code and Codex were also used as independent reviewers of drafts. The authors checked every reported value against the deposited score files by rerunning the deposited scripts (analysis_v16.py, tables_v16.py, research/VERIFY.md), checked each reference against Crossref, and are responsible for the hypotheses, interpretations and conclusions.

## Results

### Post-cutoff and self-related templates under default settings

For each protein chain, OpenFold3 retrieves a list of template chains that pass template preprocessing and supplies the first four of that list to the model. Under the unrestricted setting, at least one template released after 2021-09-30 was retrieved for 14 of 15 monomers, 16 of 16 protein–protein, 11 of 14 protein–ligand and 12 of 16 protein–nucleic-acid targets (53 of 61; 87%, 95% CI 76% to 93%) and supplied for 9, 15, 7 and 10 targets in those categories (41 of 61; 67%, 95% CI 55% to 78%). A self-related template (the target’s own entry or an entry released on the same day) was retrieved for 21 targets, 11 of them the own entry, and supplied for 16, 6 of them the own entry. All 21 were released on or before 2023-05-17. Among the 30 targets released by that date, a self-related template was supplied for 7 of 14 monomers, 3 of 5 protein–protein, 1 of 4 protein–ligand and 5 of 7 protein–nucleic-acid targets (16 of 30; 53%, 95% CI 36% to 70%; Figure 1) and retrieved but not supplied for 2, 0, 1 and 2 more (21 of 30 retrieved). For each of the 16 targets, a supplied self-related chain had 99.3% to 100% sequence identity to the target chain it was retrieved for, aligned over 94.5% to 100% of that chain (Table S6). Of the 16, 6 were supplied their own entry, 6 a same-day entry with an identical sequence over the aligned region and 4 a same-day entry at 99.3% to 99.7% identity (self_template_identity.json), so for 12 of the 16 the supplied chain carries the target’s exact sequence.

**Figure 1.**
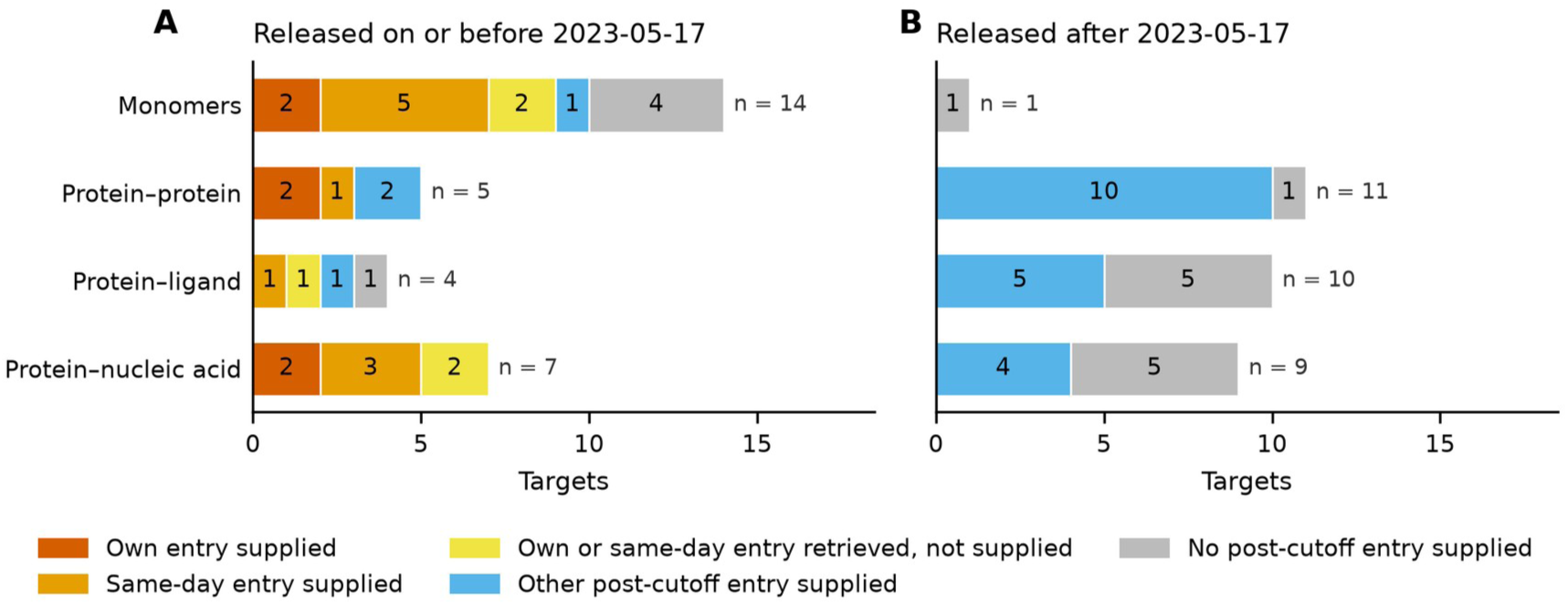
Template exposure under the unrestricted setting. Each target is classed by the closest template among the four template chains per protein chain that OpenFold3 supplies to the model: its own PDB entry, an entry released on the same day as the target, another entry released after the 2021-09-30 training cutoff, or no post-cutoff entry. Yellow marks targets whose own or a same-day entry was retrieved but ranked below the first four. (A) Targets released on or before 2023-05-17, the date of the PDB100 230517 template database, whose own entries the database could contain. (B) Targets released after 2023-05-17. Numbers in bars are target counts; n is the number of targets in each category.

**Figure 2.**
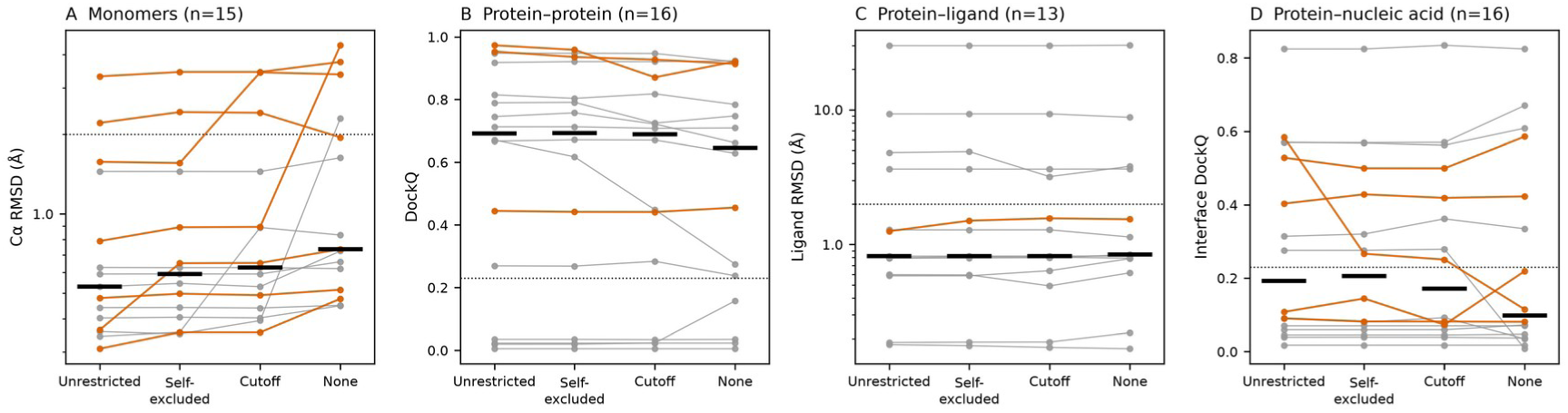
Per-target accuracy under four template settings, all run in one task per category. Each point is a target’s seed mean, the mean over three seeds of the top-ranked of five diffusion samples; lines join the same target. Orange, targets supplied their own or a same-day entry under the unrestricted setting; gray, other targets. Black bars mark medians and dotted lines the success thresholds. (A) Monomers, Cα RMSD, log scale. (B) Protein–protein complexes, DockQ. (C) Protein–ligand complexes, pocket-superposed ligand RMSD, log scale; 8CJI omitted (one unscored prediction). (D) Protein–nucleic-acid complexes, interface DockQ. Self-excluded, the unrestricted hit list without the target’s own entry and same-day entries.

**Table 1.** Highest sequence identity of target protein chains to PDB entries released on or before 2021-09-30.

| Category | Targets | Chains ≥ 90% | Targets with all chains ≥ 90% | Targets with no chain ≥ 50% |
|---|---|---|---|---|
| Monomers | 15 | 9 of 15 | 9 | 1 |
| Protein–protein | 16 | 27 of 32 | 11 | 0 |
| Protein–ligand | 14 | 13 of 14 | 13 | 0 |
| Protein–nucleic acid | 16 | 8 of 16 | 8 | 6 |

**Table 2.** Seed-mean accuracy under the four template settings (three seeds, five samples each; all settings of a category in one task on one RTX 3090). Success: Cα or ligand RMSD below 2 Å, protein–protein DockQ of at least 0.23. No success count is given for protein–nucleic-acid targets, because the DockQ thresholds have not been calibrated for nucleic-acid interfaces; their medians are reported and compared directly. Protein–ligand: 13 targets, since one 8CJI unrestricted prediction could not be scored (Materials and methods).

| Category (n) | Metric | Unrestricted | Cutoff | None | Self-excluded |
|---|---|---|---|---|---|
| Monomer (15) | median Cα RMSD (Å) | 0.53 | 0.63 | 0.73 | 0.59 |
|  | successes | 13 | 12 | 11 | 13 |
| Protein–protein (16) | median DockQ | 0.69 | 0.69 | 0.65 | 0.69 |
|  | successes | 12 | 12 | 12 | 12 |
| Protein–ligand (13) | median ligand RMSD (Å) | 0.82 | 0.82 | 0.84 | 0.82 |
|  | successes | 9 | 9 | 9 | 9 |
| Protein–nucleic acid (16) | median interface DockQ | 0.19 | 0.17 | 0.10 | 0.21 |

### Relatedness to pre-cutoff structures

Every protein–protein target had at least one chain at 90% identity or above. Six nucleic-acid-binding proteins had no pre-cutoff relative at 50% identity or above (7YUL and 8K3D had no hit at E-value 1; 7Y43, 8B4C, 8EY6 and 8U6X were between 37% and 48%). With 14 to 16 targets per category, no significant correlation between identity and seed-mean accuracy under the cutoff setting was detected (Spearman ρ between −0.35 and 0.46, all p ≥ 0.07). Identity was measured per chain and does not describe the novelty of an interface.

### Accuracy under the template settings

The accuracy comparison was sized by the targets available, not by a power calculation, so it bounds the effect of exposure rather than measuring it. Limiting templates to the cutoff shifted monomer Cα RMSD by a median −0.036 Å (unadjusted p = 0.010, Holm p = 0.11; Table 3), and one monomer (7P8G) changed success class. The rerun did not reproduce the significance of the earlier, hardware-mixed design (median −0.042 Å, Holm p = 0.010; Supplementary text S7), and the 80% detectable shift for this contrast is 0.26 Å. Only the unrestricted-versus-none monomer contrast remained significant after adjustment (−0.183 Å, Holm p = 0.031). The monomer unrestricted-versus-cutoff result is therefore inconclusive: it is not significant after adjustment and not equivalent within 0.1 Å (TOST p = 0.45). It is, however, well above run-to-run variation: for targets whose template list was identical in the unrestricted and self-excluded settings, the two differed by a median of 0.002 Å (maximum 0.016 Å) for monomers and at most 0.08 Å in any category. Without the seven development targets the monomer median was −0.041 Å (n = 14, unadjusted p = 0.005), and the three complex categories remained equivalent within their margins (TOST p = 0.039, 0.027 and 0.007). For the three complex categories, the unrestricted and cutoff settings were equivalent within the stated margins: protein–protein +0.001 DockQ (TOST p = 0.017), protein–ligand −0.002 Å (p = 0.011) and protein–nucleic acid 0.000 DockQ (p = 0.003). Success counts did not differ between the two settings in any complex category. Because templates act on protein chains, the global Cα RMSD of each complex was also compared: it was 0.154 Å lower with unrestricted templates for protein–protein complexes (unadjusted p = 0.058; interval −0.558 to +0.005 Å) and did not differ for protein–ligand (−0.000 Å) or protein–nucleic-acid complexes (−0.000 Å), so a protein-level template effect in protein–protein complexes is not excluded (extra_v16.json).

**Table 3.** Paired comparisons of per-target seed means, first setting minus second (negative favors the first for RMSD, positive for DockQ). Holm over the 12 primary tests; TOST margin 0.1 Å (monomer), 0.05 DockQ, 0.5 Å ligand RMSD; MDE = smallest median shift detected with 80% power.

| Category | Comparison | n | Median difference [interval] | Wilcoxon p | Holm p | TOST p | MDE |
|---|---|---|---|---|---|---|---|
| Monomer (Å) | Unrestricted vs cutoff | n=15 | -0.036 [-0.204, +0.000] | 0.0103 | 0.113 | 0.445 | 0.255 |
| Monomer (Å) | Cutoff vs none | n=15 | -0.067 [-0.193, +0.006] | 0.0479 | 0.443 | 0.445 | 0.609 |
| Monomer (Å) | Unrestricted vs none | n=15 | -0.183 [-0.487, -0.036] | 0.0026 | 0.031 | 0.953 | 0.809 |
| Protein–protein (DockQ) | Unrestricted vs cutoff | n=16 | +0.001 [-0.003, +0.026] | 0.1591 | 1.000 | 0.017 | 0.040 |
| Protein–protein (DockQ) | Cutoff vs none | n=16 | +0.000 [-0.014, +0.042] | 0.5619 | 1.000 | 0.004 | 0.049 |
| Protein–protein (DockQ) | Unrestricted vs none | n=16 | +0.029 [-0.000, +0.042] | 0.0443 | 0.443 | 0.012 | 0.073 |
| Protein–ligand (Å) | Unrestricted vs cutoff | n=13 | -0.002 [-0.011, +0.008] | 0.8394 | 1.000 | 0.011 | 0.162 |
| Protein–ligand (Å) | Cutoff vs none | n=14 | -0.022 [-0.206, +0.026] | 0.2958 | 1.000 | 0.000 | 0.214 |
| Protein–ligand (Å) | Unrestricted vs none | n=13 | -0.008 [-0.198, +0.149] | 0.5879 | 1.000 | 0.002 | 0.284 |
| Protein–nucleic acid (DockQ) | Unrestricted vs cutoff | n=16 | -0.000 [-0.003, +0.009] | 0.9399 | 1.000 | 0.003 | 0.020 |
| Protein–nucleic acid (DockQ) | Cutoff vs none | n=16 | -0.000 [-0.046, +0.027] | 0.8999 | 1.000 | 0.019 | 0.070 |
| Protein–nucleic acid (DockQ) | Unrestricted vs none | n=16 | -0.001 [-0.038, +0.009] | 0.4637 | 1.000 | 0.033 | 0.085 |

### Own and same-day entries

In the 7 monomers supplied their own or a same-day entry, the unrestricted setting was more accurate than the cutoff setting by a median 0.130 Å (unadjusted p = 0.016, Holm p = 0.23 within the exploratory family). Removing only the own and same-day entries gave the same median accuracy as the cutoff setting (+0.0001 Å) and matched it within 0.02 Å for 6 of the 7 (Table S11). The exception was 7P8G: its Cα RMSD was 1.58 Å unrestricted, 1.56 Å self-excluded and 3.43 Å with the cutoff, so the gain came from relatives released after the target (7PE4 on 2023-02-22 and 7PD5 on 2023-02-15, template TM-scores of 0.996 and 0.991 to the deposited structure), not from its own entry (released 2023-01-25). In the earlier self-excluded runs, which cut the template search at the day before the target’s release, these later relatives were also removed and 7P8G returned to the cutoff accuracy (Supporting information); the two designs agree once this difference is taken into account. For 7V34 the own-entry exclusion accounted for the whole difference (0.36, 0.65 and 0.65 Å). Among complexes, 7Y43 was the one target whose same-day entry carried a large change (0.58 DockQ unrestricted, 0.27 self-excluded, 0.25 with the cutoff); the other 8 exposed complexes differed between the unrestricted and cutoff settings by at most 0.11 DockQ (8BJ7, which kept 0.96 DockQ without its same-day entry) or 0.31 Å (8B9H).

Template similarity explains part of this pattern. The median TM-score of the best of the four supplied template chains to the deposited chain was 0.993, 0.989 and 0.967 for monomers under the unrestricted, self-excluded and cutoff settings, and 0.990, 0.961 and 0.864 for protein chains of protein–nucleic-acid complexes (Table S12; template_tm.py). Removing the own and same-day entries therefore left near-identical later relatives for most monomers.

Removing the MSA, with cutoff-limited templates, raised the median Cα RMSD by 0.225 Å (p = 0.005; Table 4). This is a separate contrast from the template comparisons and was not tested against them.

**Table 4.** Exploratory contrasts. The first 15 rows (targets supplied their own or a same-day entry) form one family with Holm adjustment over 15 tests; the last row (monomer MSA control) is a separate single test.

| Category | Comparison | n | Median difference [interval] | Wilcoxon p | Holm p | TOST p | MDE |
|---|---|---|---|---|---|---|---|
| Monomer (Å) | Unrestricted vs cutoff (self-supplied) | n=7 | -0.130 [-1.855, -0.012] | 0.0156 | 0.234 | 0.891 | 0.931 |
| Monomer (Å) | Cutoff vs none (self-supplied) | n=7 | -0.083 [-3.450, +0.466] | 0.3750 | 1.000 | 0.531 | 1.818 |
| Monomer (Å) | Unrestricted vs none (self-supplied) | n=7 | -0.371 [-3.554, +0.262] | 0.0781 | 0.938 | 0.922 | 1.782 |
| Monomer (Å) | Unrestricted vs self-excluded (self-supplied) | n=7 | -0.101 [-0.287, +0.017] | 0.0312 | 0.438 | 0.656 | 0.137 |
| Monomer (Å) | Self-excluded vs cutoff (self-supplied) | n=7 | +0.000 [-1.872, +0.014] | 1.0000 | 1.000 | 0.148 | 0.974 |
| Protein–protein (DockQ) | Unrestricted vs cutoff (self-supplied) | n=3 | +0.026 [n too small] | 0.2500 | 1.000 | 0.625 | – |
| Protein–protein (DockQ) | Cutoff vs none (self-supplied) | n=3 | -0.014 [n too small] | 0.5000 | 1.000 | 0.250 | – |
| Protein–protein (DockQ) | Unrestricted vs none (self-supplied) | n=3 | +0.040 [n too small] | 0.5000 | 1.000 | 0.250 | – |
| Protein–protein (DockQ) | Unrestricted vs self-excluded (self-supplied) | n=3 | +0.015 [n too small] | 0.2500 | 1.000 | 0.125 | – |
| Protein–protein (DockQ) | Self-excluded vs cutoff (self-supplied) | n=3 | +0.008 [n too small] | 0.2500 | 1.000 | 0.250 | – |
| Protein–nucleic acid (DockQ) | Unrestricted vs cutoff (self-supplied) | n=5 | +0.029 [n too small] | 0.1875 | 1.000 | 0.312 | – |
| Protein–nucleic acid (DockQ) | Cutoff vs none (self-supplied) | n=5 | -0.004 [n too small] | 0.6250 | 1.000 | 0.312 | – |
| Protein–nucleic acid (DockQ) | Unrestricted vs none (self-supplied) | n=5 | -0.020 [n too small] | 0.8125 | 1.000 | 0.312 | – |
| Protein–nucleic acid (DockQ) | Unrestricted vs self-excluded (self-supplied) | n=5 | +0.008 [n too small] | 0.8125 | 1.000 | 0.312 | – |
| Protein–nucleic acid (DockQ) | Self-excluded vs cutoff (self-supplied) | n=5 | +0.010 [n too small] | 0.0625 | 0.812 | 0.062 | – |
| Monomer (Å) | Cutoff vs cutoff without MSA | n=15 | -0.225 [-1.176, -0.025] | 0.0054 | 0.005 | 0.953 | 0.994 |

### Variation between seeds

Seed-to-seed spread was small for most targets, but a single seed could change a target’s success class (Table S3; Figure S1, Supplementary text S4). Within each category all settings shared seeds, GPU and inputs, so seed variation enters each paired contrast equally.

## Discussion

Default OpenFold3 inference supplied the target’s own entry (6 targets) or a same-day entry (10 targets) to the model for 16 of the 30 targets that its template database could contain; for each, a supplied self-related chain had at least 99.3% identity to the target chain. When every setting ran from the same inputs on the same hardware, this exposure did not produce a monomer accuracy shift that survived correction, although the comparison could not exclude shifts below about 0.26 Å, and interface accuracy in complexes was equivalent within the stated margins. The exposure is therefore frequent, and its average effect on this target set was not detected. It is concentrated in a few targets (7P8G, 7V34, 7Y43), and in the largest of them the gain came from later relatives rather than from the own entry.

The template database was inferred to be the ColabFold server’s PDB100 230517 from the server’s published history and the latest observed template release date. OpenFold3 supplies templates per protein chain, so a self-related template does not carry the placement of partner chains, ligands or nucleic acids; the complex comparisons test whether a self-related template improves the protein chains of a complex, and their equivalence says nothing about partner placement. Runs N’ Poses reported no ligand difference for AlphaFold 3 with and without templates [14]. A date-based test set built from later entries avoids own-entry retrieval but not post-cutoff relatives: 24 of the 31 later targets retrieved a post-cutoff template.

Release after the training cutoff also does not establish novelty, since close pre-cutoff sequence relatives were common.

That default template search can return a target’s own entry is an expected property of searching a database that contains it; the contribution here is the measured frequency for OpenFold3, an audit that reports it per template chain, and a controlled estimate of its effect. For a benchmark of this size, the effect of exposure on aggregate accuracy was below what the comparison can detect, but individual targets can change success class, so per-target results from date-based benchmarks remain sensitive to it.

The analysis evolved, and the accuracy results should be weighed with that in mind. An earlier design, in which the settings ran as separate jobs on mixed GPU models and pipeline revisions, found a monomer shift that survived correction (−0.042 Å, Holm p = 0.010); the single-task rerun reported here, which removes those confounds, did not (−0.036 Å, Holm p = 0.11). The equivalence margins were fixed before the rerun but after the earlier runs had been analysed, and the self-excluded setting, the template TM-scores and the MSA control were added after the earlier runs were reviewed. Each choice is described in Materials and methods, but together they make this a flexible analysis, so the accuracy comparison is reported as bounds and not as a confirmatory test. The earlier design is kept in S1 File for the record. The frequency result does not depend on these choices: it is read from the template lists the server returned.

The evidence covers one open reproduction, OpenFold3 0.4.2 with the of3-p2-155k checkpoint, under one inference configuration. The frequency describes that configuration: it is set by the template database of the server, not by the model. Predictors that obtain templates from the same server would receive the same hits, but whether their accuracy depends on them was not measured, so the accuracy bounds apply to OpenFold3 only.

### Limitations

With 13 to 16 targets per category, the monomer contrast could detect shifts of about 0.26 Å with 80% power, and the self-exposed subsets (1 to 7 targets per category) are exploratory. The equivalence bounds cover all targets of a category, most of which could not retrieve their own entry, so they do not bound the effect on the exposed targets alone. The sampling frame is weighted toward 2023 (Materials and methods), and the exposure rates describe this filtered target set, not post-cutoff PDB entries in general. The selection procedure cannot be repeated: re-running it now selects a different sample, because the query window has changed and the original candidate lists were not kept. The selected set and the selection program are archived and every target is verified against them, so the results can be reproduced, but the sample cannot be re-derived from the stated procedure; for a study of benchmark practice this is a real gap. Seven targets were also used during pipeline development; results without them are reported for every category. Only OpenFold3 was tested, on X-ray structures at 2.5 Å or better with at most 600 residues, single non-covalent ligands of 250 to 700 Da and no modified residues.

The template database version was inferred; the raw server responses of the reported runs are archived, but the server itself was not controlled and may now return different hits. MSAs came from post-cutoff databases; the MSA-free control was run for monomers with cutoff templates only, so MSA leakage of target information was not isolated. Monomers and complexes ran on two RTX 3090 devices; within a category every setting ran on one device. One protein–ligand prediction (8CJI, one seed, unrestricted) could not be scored because RDKit could not read the predicted ligand, and 8CJI is excluded from contrasts with that setting. Relatedness was measured as local sequence identity per chain, which does not establish training-set membership. Template exposure was established from the template chains passed to the model, not from how strongly the model used them.

## Conclusions

Under its default inference configuration, with templates from the ColabFold server, OpenFold3 0.4.2 received a nominally held-out target’s own entry, or a near-identical same-day entry, as a template for 16 of the 30 targets whose own entries the server’s template database could contain. With every setting run from the same inputs on one GPU, the accuracy comparison could not detect an effect of this exposure: limiting templates to the training cutoff did not change monomer accuracy detectably after correction, with 15 targets able to detect shifts of about 0.26 Å, and interface accuracy in complexes was equivalent within 0.05 DockQ or 0.5 Å. Date-based benchmarks that use inference-time templates should limit templates to the training cutoff, archive the server responses and report the template and sequence database versions, any own-entry or same-day templates supplied, target relatedness to earlier structures and the spread over several seeds.

## Data availability

All data and code are deposited at Zenodo [16], doi:10.5281/zenodo.24517157 (data CC-BY-4.0, code MIT): the target lists and the archived original selection records with the verify_cohort.py check; the OpenFold3 query files; the Nextflow prediction pipeline and a container build recipe from public inputs; for the single-task runs of the main text, every predicted model with its confidence file, the template chains each setting kept, the release dates, the archived ColabFold MSA and template-hit responses, the GPU records and the score files; the data of the earlier runs reported in the Supporting information; and the target-selection, scoring, statistics, table and figure scripts. OpenFold3 0.4.2 and its checkpoint are public [2]; the container digest and non-default settings are given in Materials and methods. The container image itself is held on the authors’ internal registry and is rebuilt from the deposited recipe. Reference structures are available from the Protein Data Bank [15] under the identifiers in Table S1.

## Author contributions

Conceptualization: Gabriel Richman, Mahdi Soultana. Investigation, Software, Formal analysis, Visualization: Mahdi Soultana. Validation: Olamide Isreal Odesanya, Mervin Gaitho. Supervision: Gabriel Richman. Writing – review and editing: Mahdi Soultana, Olamide Isreal Odesanya, Mervin Gaitho, Gabriel Richman. (To be confirmed by the authors.)

## Acknowledgments

We thank the RCSB PDB for public structure data and the ColabFold team for the public MSA and template server. The study was carried out on the Omic research platform (https://omic.ai); the use of AI tools is described in Materials and methods.

## Supporting information

- Table S1 (table_S1_v5.tsv): all 61 targets with release date, resolution, ligand, pre-cutoff identity, own-entry, same-day and post-cutoff template counts.
- Table S2 (table_S2_gpu.tsv): GPU model per run.
- Table S3 (table_S3_seeds.tsv): per-seed top-ranked values.
- Table S4 (table_S4_posebusters.tsv): PoseBusters redock validity for the top-ranked ligand prediction per target, setting, and seed; the reference-pose RMSD criterion is excluded from the validity flag.
- Table S5 (matched_gpu_controls.tsv, matched_gpu_controls.json): per-target same-model-GPU control values for Table S7.
- Table S6 (template_chains.json): for each target protein chain, the retrieved template chains in order, which were supplied to the model, their release dates, and the identity and coverage of own-entry and same-day chains.
- Table S10 (scores_v3/table_S10_selfx_arm.md, with inputs/selfx/ the per-release query files and scores_v3/selfx/ the cluster scores of the self-excluded arm).
- Table S11 and Table S12 (below; numbers_v16.json, tm_.tsv): per-target values of the 16 targets supplied their own or a same-day entry, and TM-score of the supplied template chains to the deposited chains, from the single-task runs of the main text.
- v16/: the single-task runs of the main text: per-category arms_status.tsv, templates_by_arm.tsv (template chains each setting kept), release_dates.json, gpu.txt, score files scores/-_scores.csv, and analysis_v16.py, tables_v16.py, template_tm.py, fig2_v16.py.
- Tables S2, S3 and S5 to S10 and Supplementary text S2 to S8 and both arm sections that follow them describe EARLIER RUNS, superseded by the single-task runs of the main text. In those runs each template setting ran as a separate job on varying GPU models and pipeline revisions, with MSAs fetched again for each job, and the self-excluded setting cut the template search at the day before the target’s release, which also removed relatives released after the target. They are kept so readers can compare the two designs; where their interpretation differs from the main text (for example 7P8G, Table S10), the main text supersedes them. Every number in the main text comes from the single-task runs.
- Supplementary text S1 to S8 and Figure S1 (below): material moved from the main text (selection funnel, GPU models, software and scoring provenance, per-target and seed detail, ligand validity and failures, release-date strata, robustness), then the self-excluded and no-template arm text with Tables S9 and S10.
- software_versions.json and select_funnel.json: software versions and run settings, and selection-funnel counts of the later re-run.
- replicate_noise.json, scores_replicate/ and gpu_per_run.tsv: repeated-prediction differences for the shared seed and the GPU device of each monomer run; self_template_identity.json: own, identical same-day and near-identical same-day templates per target.
- openfold3_pipeline/: the Nextflow prediction pipeline, its configuration and the container build recipe; monomer_runs.tsv: pipeline revision and GPU device of each monomer run.
- original_selection/: the selection as made (selected_set.json, pilot_targets_manifest.json, pilot_candidates_rcsb_detail.json, seed_seqs.json, select_set.py) and verify_cohort.py, which checks every target of this paper against it.

**Table S11.** Seed-mean values of the targets supplied their own or a same-day entry under the unrestricted setting (Cα RMSD or ligand RMSD in Å, DockQ for interfaces).

| Category | Target | Unrestricted | Self-excluded | Cutoff | None |
|---|---|---|---|---|---|
| Monomer | 7P8G | 1.575 | 1.558 | 3.430 | 3.369 |
| Monomer | 7SJV | 0.309 | 0.356 | 0.356 | 0.477 |
| Monomer | 7V34 | 0.363 | 0.650 | 0.652 | 0.735 |
| Monomer | 7XYI | 0.789 | 0.890 | 0.893 | 4.343 |
| Monomer | 7XZ4 | 3.311 | 3.441 | 3.441 | 3.757 |
| Monomer | 7ZGD | 2.207 | 2.425 | 2.411 | 1.946 |
| Monomer | 8DJX | 0.480 | 0.498 | 0.492 | 0.516 |
| Protein–protein | 8AY8 | 0.953 | 0.936 | 0.927 | 0.914 |
| Protein–protein | 8BJ7 | 0.973 | 0.959 | 0.871 | 0.921 |
| Protein–protein | 8CWR | 0.445 | 0.442 | 0.441 | 0.455 |
| Protein–ligand | 8B9H | 1.255 | 1.503 | 1.565 | 1.539 |
| Protein–nucleic acid | 7MXV | 0.528 | 0.499 | 0.499 | 0.586 |
| Protein–nucleic acid | 7WKP | 0.403 | 0.428 | 0.419 | 0.423 |
| Protein–nucleic acid | 7Y43 | 0.584 | 0.267 | 0.251 | 0.115 |
| Protein–nucleic acid | 7YUL | 0.090 | 0.082 | 0.082 | 0.081 |
| Protein–nucleic acid | 8EDJ | 0.108 | 0.144 | 0.073 | 0.219 |

**Table S12.**
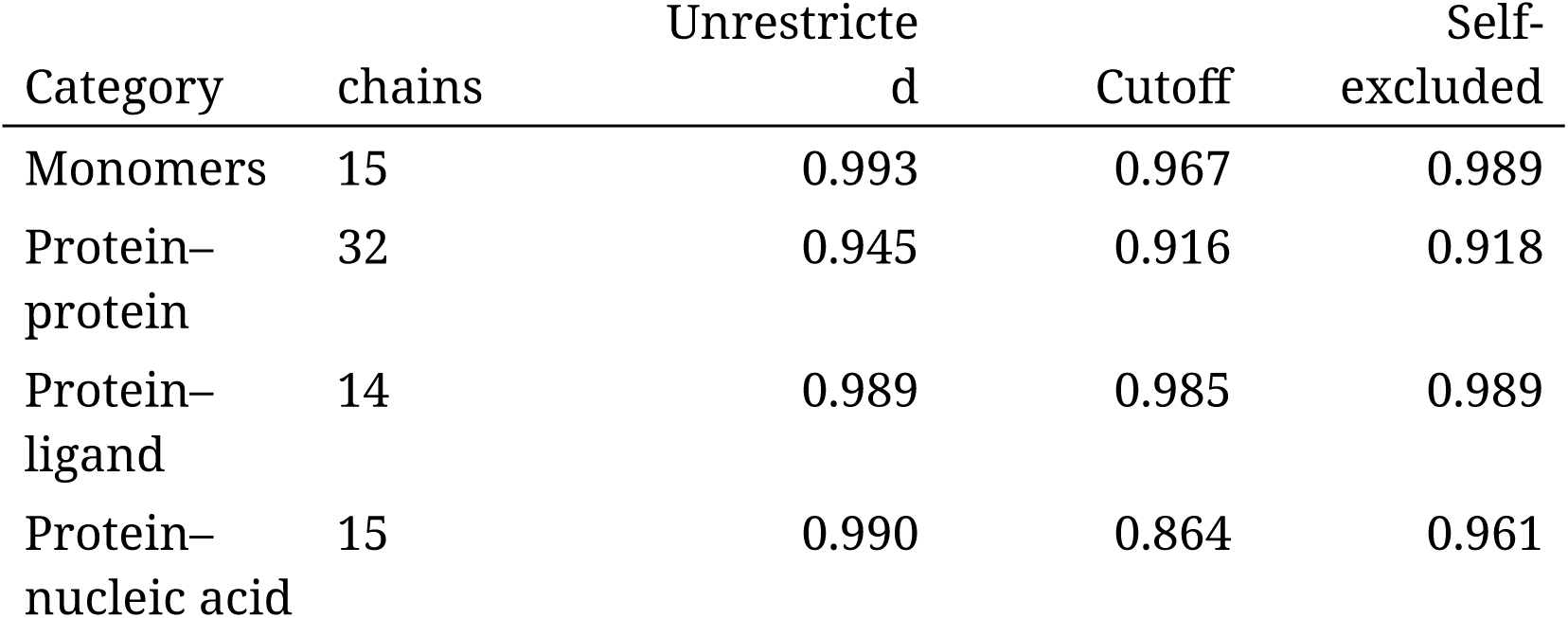
Median TM-score, normalized by the deposited chain, of the best of the four template chains supplied for each protein chain. Median TM-score (reference-normalised) of the best of the four supplied template chains.

| Category | chains | Unrestricted | Cutoff | Self-excluded |
|---|---|---|---|---|
| Monomers | 15 | 0.993 | 0.967 | 0.989 |
| Protein–protein | 32 | 0.945 | 0.916 | 0.918 |
| Protein–ligand | 14 | 0.989 | 0.985 | 0.989 |
| Protein–nucleic acid | 15 | 0.990 | 0.864 | 0.961 |

**Table S7.**
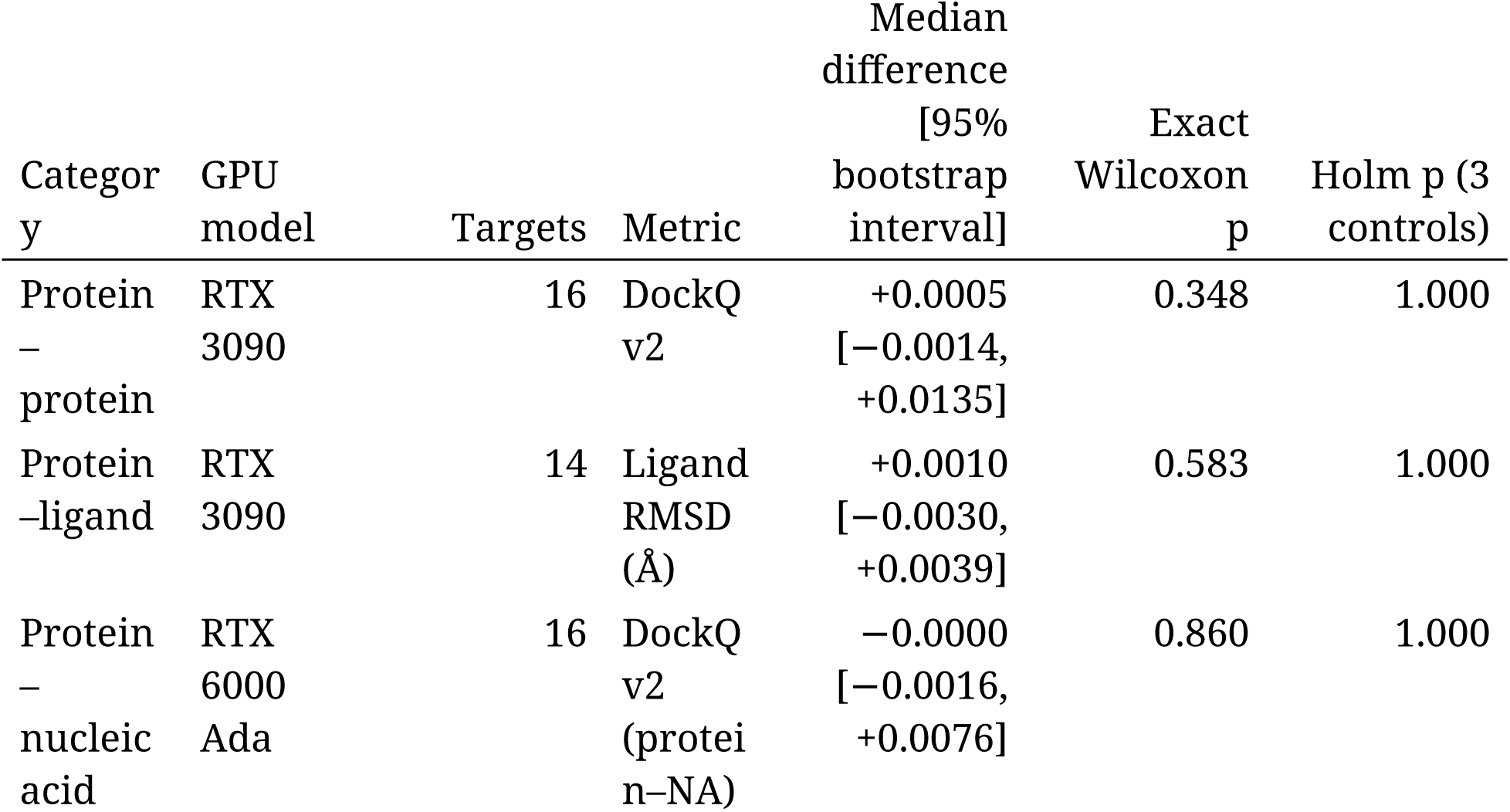
Unrestricted-minus-cutoff control differences (earlier runs) (per-target values in Table S5) using three seeds on the same GPU model for each category. Each target contributes the mean of its top-ranked prediction across the same three seed identifiers; intervals are bootstrap percentile intervals for the median paired difference.

| Category | GPU model | Targets | Metric | Median difference [95% bootstrap interval] | Exact Wilcoxon p | Holm p (3 controls) |
|---|---|---|---|---|---|---|
| Protein–protein | RTX 3090 | 16 | DockQ v2 | +0.0005 [−0.0014, +0.0135] | 0.348 | 1.000 |
| Protein–ligand | RTX 3090 | 14 | Ligand RMSD (Å) | +0.0010 [−0.0030, +0.0039] | 0.583 | 1.000 |
| Protein–nucleic acid | RTX 6000 Ada | 16 | DockQ v2 (protein–NA) | −0.0000 [−0.0016, +0.0076] | 0.860 | 1.000 |

**Table S8.** Spread of the top-ranked metric across three seeds. SD is computed across the three seeds for each target, then summarized by the median across targets.

| Category | Median target-level SD (unrestricted / cutoff / none) | Largest target range (target, setting) |
|---|---|---|
| Monomers (Å) | 0.016 / 0.033 / 0.036 | 2.95 (7ZO6, none) |
| Protein–protein (DockQ v2) | 0.008 / 0.017 / 0.016 | 0.51 (8J64, none) |
| Protein–ligand (Å) | 0.057 / 0.065 / 0.054 | 7.08 (8FRJ, none) |
| Protein–nucleic acid (DockQ v2) | 0.008 / 0.010 / 0.009 | 0.81 (7U6K, cutoff) |

### Supplementary text: methods and results detail moved from the main text

*S1 describes the target selection used throughout. S2 to S8 and the self-excluded and no-template arm sections report the EARLIER design, in which the settings ran as separate jobs on mixed GPU models and pipeline revisions. That design is superseded by the single-task rerun in the main text and is kept for the analysis history described in the Discussion. Where a number here differs from the main text, the main text is current*.

#### S1 Target-selection funnel

The selection as made is archived in data/raw/original_selection/. selected_set.json holds the 56 systematically selected entries, 14 per category, with release date, resolution and title; pilot_targets_manifest.json holds the seven targets also used for pipeline development, with the RCSB detail response they were taken from; seed_seqs.json holds the nine development sequences excluded by the identity filter. verify_cohort.py accounts for all 61 targets of this paper, 54 systematically selected and 7 development targets, and fails if any is unaccounted for. Of the 56 systematically selected entries, 54 were predicted; 8AAE and 8GAR were excluded for the reasons given in Materials and methods.

Re-running the same queries, seed and rules against the RCSB at the time of writing examined 16, 94, 89 and 27 candidates to fill 14 places in the monomer, protein–protein, ligand and protein–nucleic-acid categories; identity of 30% or more to an accepted or development chain rejected 1, 71, 53 and 9 of them, non-standard residues 1, 6, 2 and 3, and the category rules 0, 1, 20 and 1 (protein– protein candidates with similar chains rejected 2 more). That re-run selected 0, 0, 1 and 4 of the published targets (select_funnel.json). Because the program takes the first 400 entries ordered by release date before shuffling, a change in the content of the window changes the candidate list and therefore the sample, so these counts describe the procedure on the later list. The candidate lists returned by the original queries were not retained.

#### S2 GPU models of the runs

Each target and setting was predicted with three seeds and five diffusion samples on RTX-series GPUs; the GPU model of each run is listed in Table S2. Both monomer settings in the primary contrast ran on an RTX 3090, and the monomer no-template arm of the original runs on an RTX 6000 Ada, so contrasts involving that arm are GPU-confounded in Tables 2 and 3; the monomer, protein–ligand and protein–protein no-template arms were repeated on an RTX 3090 (Table S9), while the protein–nucleic-acid no-template arm could not be obtained on an RTX 6000 Ada and was run on an RTX 3090. In the primary three-seed interface runs the unrestricted and cutoff arms ran on different GPU models. The same-model-GPU comparison repeated the unrestricted arm with the same seed identifiers on the GPU model of each cutoff arm: RTX 3090 for protein–protein and ligand (on a different physical device) and RTX 6000 Ada for protein–nucleic acid (on the same physical device). Statistics use the three-seed runs and the same-model-GPU runs. The ligand centroid distances used a separate set of single-seed predictions (four of them on a GTX 1070), whose values are listed in Table S1.

The same-model-GPU controls (Table S7; per-target values in Table S5) compare unrestricted with cutoff-limited templates on one GPU model per category, with three seeds each. None of the three controls showed a detectable category-level shift. With the unrestricted arm on the cutoff arm’s GPU model, 7Y43 and 8BJ7 differed from cutoff by +0.341 and +0.108 DockQ v2, close to the original differences. The control compares the two template settings on one GPU model; the cutoff arm itself was not repeated. For 8BJ7 the difference rests on one cutoff seed (0.734 against 0.931 and 0.929 for the other two).

#### S3 Software versions and scoring provenance

The no-template arms of Table S9 were scored on the cluster by the of3_score tool, which runs the paper’s score_top.py and dockq_v2_all.py unchanged on the top-ranked model of each seed; the interface scores of the original runs (Tables 2 and 3) are DockQ v2 values from the same dockq_v2_all.py (Supplementary text S3).

Models were compared with the first model of the deposited structure using label sequence numbering. For each seed, the sample ranked first by OpenFold3’s sample ranking score was scored, and per-target values are the mean over three seeds. This one definition is used in every table, figure and number of the paper (Tables 2z 3, S1, S3 and S7 to S10). An earlier in-house reimplementation of the DockQ formula is superseded and not reported (Supplementary text S3).

The container held OpenFold3 0.4.2 with torch 2.10.0, pytorch-lightning 2.6.1, cuEquivariance 0.10.0, gemmi 0.7.5 and numpy 2.3.5 (versions.txt of the run), and runs were executed with Nextflow 26.10.2 (WES run records). Scoring used biopython 1.88, DockQ 2.1.3, gemmi 0.7.5, PoseBusters 0.6.5, RDKit 2026.9.2, tmtools 0.3.0, numpy 1.26.4, scipy 1.17.1 and statsmodels 0.15.0, pinned in requirements-v14.txt. Versions and run settings are in software_versions.json.

The no-template arms of Table S9 were scored on the cluster by the of3_score tool (Nextflow, one task per target, python:3.12 with the pinned packages below; code in code/upstream/of3_score/), which runs the paper’s score_top.py and dockq_v2_all.py unchanged on the top-ranked model of each seed. The interface scores of the original runs (Tables 2 and 3) are DockQ v2 values from the same dockq_v2_all.py, computed in an earlier pass on this host from the downloaded top models and stored in scores_v2/dockq_v2.json; they were not recomputed through of3_score, which cannot read those run folders. analysis_v15_original_runs.py writes numbers_v15_original_runs.json from them with the statistics of analysis_v14.py.

Models were compared with the first model of the deposited structure using label sequence numbering. For each seed, the sample ranked first by OpenFold3’s sample ranking score was scored, and per-target values are the mean over three seeds.

All DockQ values in this paper come from DockQ v2 (2.1.3); an earlier in-house reimplementation of the formula is not reported, and its scores remain only in scores/ for the record (for protein–nucleic acid it gave a lower median than DockQ v2, 0.14 against 0.24 unrestricted, so the two must not be mixed). DockQ thresholds (0.23 acceptable, 0.49 medium) were defined for protein–protein interfaces and are applied unchanged to protein–nucleic-acid interfaces.

#### S4 Monomer, success-count and seed detail

When a target counted as successful only if all three seeds met the threshold, the success counts (unrestricted, cutoff, none) were 13, 12 and 10 for monomers; 11, 11 and 10 for protein–protein; 9, 9 and 9 for protein–ligand; and 6, 5 and 5 for protein–nucleic-acid targets.

The largest monomer changes were in 7P8G (−1.856 Å; own entry supplied), 8AAQ (−0.545 Å; self-related template retrieved but not supplied) and 7V34 (−0.288 Å; same-day entries supplied). Own-entry retrieval did not always change the prediction: 7TUO retrieved its own entry, ranked below the four supplied templates, and changed by less than 0.001 Å.

The largest per-target interface changes were 7Y43 (+0.339 DockQ v2, same-day template; cutoff seeds 0.53, 0.11 and 0.12), 8J64 (+0.139, no self-related template) and 8BJ7 (+0.109, own entry). For 8J64 and for 8FRJ (+1.682 Å ligand RMSD, worse with unrestricted templates) the seed ranges of the two settings overlapped (Table S3).

Seed-to-seed spread was small for most targets (Table S8). Nine targets changed success class between seeds of at least one setting: the monomers 7ZGD (all three settings) and 7ZO6 (no templates), the protein–protein targets 8GZV (all three settings), 8J64 and 8EZL (no templates), the protein–nucleic-acid targets 7U6K (unrestricted and cutoff), 8EDJ (unrestricted and none) and 7Y43 (cutoff), and the ligand target 8FRJ (no templates). For example, 8J64 ranged from 0.02 to 0.53 DockQ, 8EZL from 0.02 to 0.44, 7Y43 from 0.11 to 0.53 (cutoff), and 8FRJ from 0.98 to 8.06 Å. A single seed can therefore change a target’s success class (Table S3, Figure 3).

**Figure S1.**
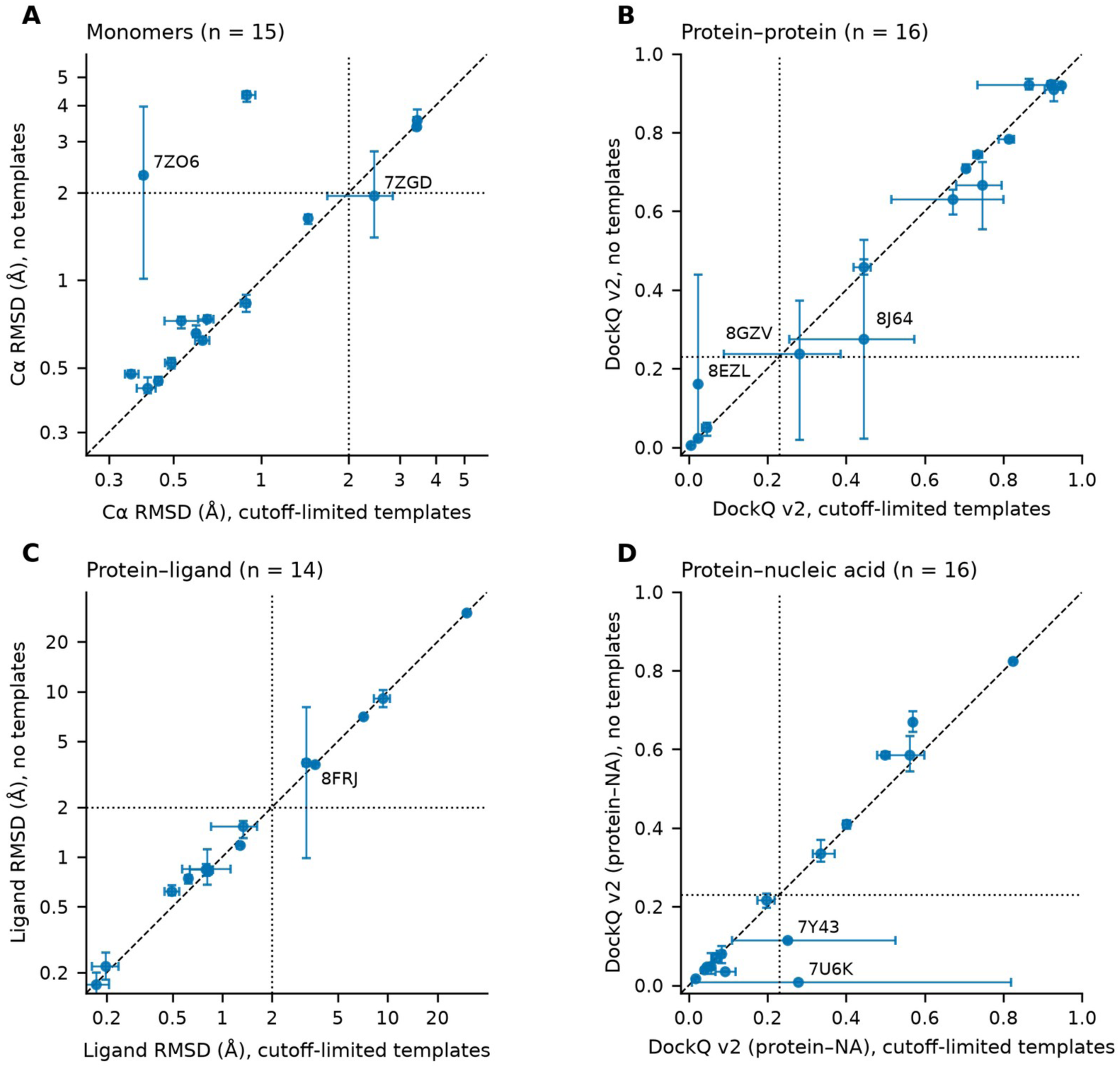
Seed-to-seed variation under cutoff-limited templates and no templates. Points are means over three seeds of the top-ranked prediction; error bars span the minimum and maximum seed values. The dashed line marks equality and dotted lines mark the success thresholds; an error bar that crosses a dotted line belongs to a target whose success class differs between seeds. Labeled targets are those named in the text. (A) Monomers, Cα RMSD, log scales. (B) Protein–protein complexes, DockQ. (C) Protein–ligand complexes, ligand RMSD, log scales. (D) Protein–nucleic-acid complexes, interface DockQ.

#### S5 Ligand pose validity and interface failures

Among the top-ranked ligand predictions, PoseBusters validity was 36/42 for unrestricted templates, 36/42 for cutoff-limited templates, and 35/42 with no templates (14 targets × three seeds per setting). Seventeen of the 19 failures were tetrahedral-chirality failures in two ligands, 8E4A (9 of 9) and 8RU8 (8 of 9), across all three settings; the other two were a chirality failure in 8BSN (no templates) and a minimum-distance-to-protein failure in 8CJI (unrestricted). The input SMILES of 8E4A and 8RU8 assign the same configuration to both stereocenters as the deposited ligands, and the failures occurred under all three template settings. Validity counts refer to the selected pose and are not a measure of pose accuracy (Table S4).

The median pocket Cα RMSD of the ligand predictions was 0.7 Å in each setting. Of the five ligands above 2 Å under the cutoff setting, 8CRC was placed in a different site (centroid 29.8 Å from the crystal position), 8BSN and 8CJI were displaced by 5.0 and 3.1 Å, 8BM8 was in the correct site (centroid 0.2 Å) with a 3.6 Å pose error in every sample and seed, and 8FRJ is a fusion construct whose domains were misplaced relative to each other (global Cα RMSD 14 Å). Protein– nucleic-acid interfaces had the lowest DockQ values of the interface categories, with 8 to 10 of 16 below 0.23 (seed mean) in each setting, a threshold not calibrated for nucleic-acid interfaces. In 8B4C the DNA duplex was predicted accurately (DNA–DNA DockQ 0.84) while the protein was not placed on it (protein –DNA DockQ 0.018; RMSD of the protein after superposition on the DNA about 40 Å) in every setting and seed.

#### S6 Interface results by template-database date

Among targets released on or before 2023-05-17, the median unrestricted-minus-cutoff difference in the original runs was +0.008 DockQ v2 for protein–protein (n = 5, unadjusted p = 0.31), +0.005 Å ligand RMSD (n = 4, p = 0.125; positive is worse) and +0.008 DockQ v2 for protein–nucleic acid (n = 7, p = 0.22). In the same-model-GPU control runs the corresponding medians were +0.001 DockQ v2 (4 of 5 targets higher), +0.002 Å (3 of 4 worse) and +0.010 DockQ v2 (5 of 7 higher). Among later targets the original medians were −0.0002 DockQ v2 (n = 11), +0.0002 Å (n = 10) and 0.000 DockQ v2 (n = 9). None of the original-run tests was significant, and with 4 to 7 targets per stratum these tests have little power; the medians are reported for completeness and are not evidence of a shift.

#### S7 Robustness and sensitivity analyses

As a sensitivity analysis that does not select the top-ranked sample, each target’s value was also averaged over all 15 outputs (three seeds by five diffusion samples); the target remains the unit of inference. Analyses excluding development targets and targets with undated cutoff-arm templates, and analyses stratified by release on or before versus after 2023-05-17, were computed from the same score files.

Excluding the one monomer development target (8T7K) left the monomer unrestricted-versus-cutoff median difference at −0.043 Å (n = 14; unadjusted p = 0.0017). Excluding 7ZO6, 8BJ7 and 8U6X, whose cutoff-arm template entries lack release dates, left median differences of −0.030 Å for monomers (n = 14), +0.0001 DockQ v2 for protein–protein (n = 15), +0.001 Å for ligand RMSD (n = 14), and 0.000 DockQ v2 for protein–nucleic acid (n = 15). Averaging all 15 outputs gave a ligand-RMSD median difference within 0.001 Å of zero; DockQ v2 was computed for the top-ranked model of each seed only, so the all-output sensitivity analysis is not reported for interfaces.

### S8 Controls, interface failures and database-date strata (moved from the main text)

### Interface results by template-database date

Among targets released on or before 2023-05-17 the unrestricted-minus-cutoff interface and ligand medians were within 0.008 of zero (DockQ v2 or Å; n = 4 to 7 per category) and none of the tests was significant; with so few targets per stratum the medians are not evidence of a shift (Supplementary text S6).

### Same-model-GPU control runs

With the unrestricted arm repeated on the cutoff arm’s GPU model, none of the three controls showed a detectable category-level shift (Table S7; per-target values in Table S5; Supplementary text S2). The cutoff arm itself was not repeated.

### No-template arms on one GPU model

The monomer, protein–protein and protein–ligand no-template arms were repeated on an RTX 3090, the GPU model of the other arms of those categories, and scored with DockQ v2 (Table S9 in the Supplementary information). The monomer result was unchanged (unrestricted minus cutoff −0.042 Å, Holm p = 0.010; unrestricted against none −0.186 Å, Holm p = 0.037); no contrast was detectable for protein–protein or protein–ligand targets. The protein–nucleic-acid no-template arm could not be run on an RTX 6000 Ada and ran on an RTX 3090, so its contrasts with the other two arms are GPU-confounded.

### Interface failures

Pose accuracy and validity failures concentrated in a few targets: of the five ligands above 2 Å under the cutoff setting, 8CRC was placed in a different site, and 8 to 10 of 16 protein–nucleic-acid targets were below DockQ 0.23 (seed mean) in each setting, a threshold not calibrated for nucleic-acid interfaces. PoseBusters validity was 36/42, 36/42 and 35/42 for unrestricted, cutoff and no templates (Supplementary text S5; Table S4).

### Robustness

Excluding the one monomer development target (8T7K) left the monomer unrestricted-versus-cutoff median difference at −0.043 Å (n = 14; unadjusted p = 0.0017); excluding the three targets with undated cutoff-arm templates left it at −0.030 Å (n = 14). The other categories were unchanged (Supplementary text S7).

## Supplementary text: self-excluded and no-template arms

### Self-excluded template arm

The self-excluded arm (Materials and methods, setting 4) was run for all 21 targets with a retrieved own or same-day template (three seeds, five samples; predictions on the cluster, runs listed in inputs/selfx/runs.json, scored with of3_score and DockQ v2). All 21 targets were scored with the same tool and commit. Interface arms ran on an RTX 3090, except that the protein–nucleic-acid unrestricted and cutoff arms ran on an RTX 6000 Ada, so protein–nucleic-acid contrasts with this arm are GPU-confounded (the 6000 Ada was unavailable for over an hour and the arm was resubmitted on a 3090). Per-target values are in Table S10. The comparisons are descriptive and unadjusted; the Holm family of Table 3 was not extended.

Among the 9 monomers the median self-excluded-minus-cutoff difference was +0.000 Å (p = 0.73; this does not establish equivalence) and was worse than the unrestricted arm for all 9 (median U minus X −0.112 Å, exact two-sided Wilcoxon p = 0.0039, the smallest p attainable with 9 targets). This p value is unadjusted: it is not part of the 12-test Holm family of Table 3, and it is one of eight unadjusted tests of Table S10 (four categories, two contrasts each); Holm-adjusted across those eight tests it is 0.031. The arm was added after the primary analysis, is exploratory, and no adjustment was pre-specified. The two monomers whose own entry was retrieved but not supplied (7TUO, 8AAQ) changed by about 0.01 Å; the larger changes were in targets supplied an own or same-day template (7P8G −1.82 Å, 7V34 −0.29 Å, 7ZGD −0.24 Å). For 8AAQ the self-excluded value matched the unrestricted value and not the cutoff value (0.354, 0.344 and 0.888 Å), so templates released between 2021-09-30 and its release date, not its own entry, carried that target’s difference. For the interface and ligand targets the self-excluded values did not differ detectably from either arm: median U minus X +0.013 DockQ v2 (n = 3, protein–protein), +0.011 (n = 7, protein–nucleic acid) and +0.048 Å (n = 2, ligand). The one large interface change, 7Y43 (+0.341 DockQ v2 unrestricted over self-excluded), was removed entirely by excluding its same-day entry (self-excluded 0.251, cutoff 0.250). In that earlier design this arm did not change the monomer estimate (−0.042 Å, Holm p = 0.0103); the arm supports the interpretation that it comes from own and same-day entries, within the limits above (nine monomers, three targets with extra later templates removed, no control for MSA information).

**Table S10.** Self-excluded template arm, per target (seed means over three seeds), and category medians. U, unrestricted; X, self-excluded; C, cutoff; differences are first minus second (negative favors the first setting for the RMSD categories, positive for DockQ v2). Medians are over the targets of this arm only; p values are exact two-sided Wilcoxon signed-rank tests, unadjusted and outside the Holm family of Table 3. Protein–nucleic-acid rows: the self-excluded arm ran on an RTX 3090 and the unrestricted and cutoff arms on an RTX 6000 Ada, so the protein– nucleic-acid contrasts in this table are GPU-confounded.

| Category | Target | Own or same-day template supplied | Unrestricted (U) | Self-excluded (X) | Cutoff (C) | U − X | X − C |
|---|---|---|---|---|---|---|---|
| Monomer, Cα RMSD (Å) | 7P8G | yes | 1.575 | 3.390 | 3.430 | -1.816 | -0.040 |
| Monomer, Cα RMSD (Å) | 7SJV | yes | 0.312 | 0.356 | 0.356 | -0.044 | +0.000 |
| Monomer, Cα RMSD (Å) | 7TUO | no | 1.447 | 1.448 | 1.447 | -0.001 | +0.001 |
| Monomer, Cα RMSD (Å) | 7V34 | yes | 0.363 | 0.656 | 0.651 | -0.292 | +0.005 |
| Monomer, Cα RMSD (Å) | 7XYI | yes | 0.788 | 0.900 | 0.892 | -0.112 | +0.008 |
| Monomer, Cα RMSD (Å) | 7XZ4 | yes | 3.313 | 3.445 | 3.440 | -0.132 | +0.005 |
| Monomer, Cα RMSD (Å) | 7ZGD | yes | 2.201 | 2.436 | 2.446 | -0.235 | -0.010 |
| Monomer, Cα RMSD (Å) | 8AAQ | no | 0.344 | 0.354 | 0.888 | -0.011 | -0.534 |
| Monomer, Cα RMSD (Å) | 8DJX | yes | 0.474 | 0.491 | 0.491 | -0.017 | -0.000 |
| Protein–protein, DockQ v2 | 8AY8 | yes | 0.954 | 0.936 | 0.928 | +0.018 | +0.008 |
| Protein–protein, DockQ v2 | 8BJ7 | yes | 0.972 | 0.959 | 0.865 | +0.013 | +0.094 |
| Protein–protein, DockQ v2 | 8CWR | yes | 0.446 | 0.440 | 0.445 | +0.006 | -0.005 |
| Protein–ligand, ligand RMSD (Å) | 8B9H | yes | 1.278 | 1.331 | 1.328 | -0.053 | +0.003 |
| Protein–ligand, ligand RMSD (Å) | 8BSN | no | 9.308 | 9.159 | 9.305 | +0.148 | -0.146 |
| Protein–nucleic acid, DockQ v2 | 7MXV | yes | 0.528 | 0.499 | 0.498 | +0.029 | +0.001 |
| Protein–nucleic acid, DockQ v2 | 7U6K | no | 0.274 | 0.276 | 0.279 | -0.002 | -0.003 |
| Protein–nucleic acid, DockQ v2 | 7WKP | yes | 0.410 | 0.399 | 0.400 | +0.011 | -0.001 |
| Protein–nucleic acid, DockQ v2 | 7Y43 | yes | 0.592 | 0.251 | 0.250 | +0.341 | +0.000 |
| Protein–nucleic acid, DockQ v2 | 7YUL | yes | 0.090 | 0.086 | 0.083 | +0.005 | +0.003 |
| Protein–nucleic acid, DockQ v2 | 8DW1 | no | 0.824 | 0.835 | 0.824 | -0.010 | +0.010 |
| Protein–nucleic acid, DockQ v2 | 8EDJ | yes | 0.214 | 0.196 | 0.197 | +0.018 | -0.000 |
| Category | n | Median U − X | exact Wilcoxon p (unadjusted) | Median X − C | exact Wilcoxon p (unadjusted) |  |  |
| Monomer, Cα RMSD (Å) | 9 | -0.112 | 0.004 | +0.000 | 0.734 |  |  |
| Protein–protein, DockQ v2 | 3 | +0.013 | 0.250 | +0.008 | 0.500 |  |  |
| Protein–ligand, ligand RMSD (Å) | 2 | +0.048 | 1.000 | -0.072 | 1.000 |  |  |
| Protein–nucleic acid, DockQ v2 | 7 | +0.011 | 0.109 | +0.000 | 0.688 |  |  |

### No-template arms on one GPU model, scored with DockQ v2

The no-template arms of the original runs differed from their template arms in GPU model for monomers and interfaces. The monomer and protein–ligand no-template arms were repeated with three seeds and five samples on an RTX 3090 (monomer run cc87e992, ligand run a0f6bf79); the protein–protein no-template arm was not repeated, because in the original runs it already ran on the same RTX 3090 device as the protein–protein cutoff arm, so its cutoff-versus-none values in this table are those of Table 3. All arms were and scored on the cluster with the same scripts as the other arms (the of3_score tool; Methods) and with DockQ v2 on every interface. The protein–nucleic-acid no-template arm could not be run on an RTX 6000 Ada, the GPU model of its unrestricted and cutoff arms: two submissions (7b0808f1, 6ad5e4ce) waited for that model for 65 and 29 minutes and failed because no such GPU became free, so the arm was run on an RTX 3090 (39b0eb8f) and its contrasts with the other two arms remain GPU-confounded.

Per-target values are in scores_v3/; the table was computed by analysis_v14.py and make_table_v14_noarms.py.

**Table S9.** Seed-mean accuracy and paired contrasts involving the no-template arm, with every arm of a category on one GPU model except protein–nucleic acid. Medians are for unrestricted (U), cutoff (C) and no templates (None). Differences are the first setting minus the second; the Holm adjustment covers the same 12 contrasts as Table 3 and was recomputed from the same-GPU arms (Tables 2 and 3 use the original runs, whose arms for some categories ran on different GPU models, so the template-arm medians of this table differ slightly from theirs; interface values in both are DockQ v2). Protein–protein and protein–nucleic-acid values are DockQ v2 on all native interfaces.

| Category (n) | GPU of arms | Metric | Median U / C / None | Cutoff vs none: median difference [interval], p, Holm p | Unrestricted vs none: median difference [interval], p, Holm p |
|---|---|---|---|---|---|
| Monomer (15) | RTX 3090 | Cα RMSD (Å) | 0.531 / 0.628 / 0.736 | -0.062 [-0.186, +0.008], 0.0479, 0.479 | -0.186 [-0.488, -0.026], 0.0034, 0.037 |
| Protein–protein (16) | RTX 3090 | DockQ v2 | 0.692 / 0.687 / 0.649 | +0.000 [-0.010, +0.039], 0.4954, 1.000 | +0.025 [-0.004, +0.045], 0.0654, 0.589 |
| Protein–ligand (14) | RTX 3090 | ligand RMSD (Å) | 1.049 / 1.051 / 1.014 | -0.038 [-0.200, +0.004], 0.0785, 0.628 | -0.023 [-0.197, +0.149], 0.3575, 1.000 |
| Protein–nucleic acid (16) | RTX 6000 Ada (no-template arm: RTX 3090) | DockQ v2, protein–NA interfaces | 0.244 / 0.224 / 0.098 | -0.001 [-0.038, +0.002], 0.4637, 1.000 | -0.002 [-0.058, +0.010], 0.3484, 1.000 |

With the monomer arms on one GPU model the monomer result was the same as in Table 3: the median Cα RMSD was 0.53 Å (unrestricted), 0.63 Å (cutoff) and 0.74 Å (none), 13, 12 and 11 targets were below 2 Å, and the unrestricted-minus-cutoff difference was −0.042 Å (Holm p = 0.010; unchanged). Against no templates the unrestricted difference was −0.186 Å (Holm p = 0.037); the earlier value from the RTX 6000 Ada arm was −0.184 Å. For protein–ligand targets no contrast with the no-template arm was detectable (median ligand RMSD 1.05, 1.05 and 1.01 Å; 9 of 14 below 2 Å in each arm); PoseBusters validity of the selected poses was 37, 36 and 36 of 42 for unrestricted, cutoff and no templates (the single no-template failure outside 8E4A and 8RU8 was a chirality failure in 8BSN). For protein– protein and protein–nucleic-acid targets DockQ v2 medians were 0.69, 0.69 and 0.65, and 0.24, 0.22 and 0.10; no contrast was detectable after adjustment, and the unrestricted-minus-cutoff differences were +0.0005 and −0.0000. Success classes of three protein–protein targets (8EZL, 8GZV, 8J64) and one protein–nucleic-acid target (8EDJ) changed between seeds of the no-template arm. DockQ v2 was computed on all interfaces of every model; the monomer DockQ step reported an error for the three seeds of 8B2C (chain identifier AAA exceeds the PDB-format limit), which does not enter any monomer value. Interface LDDT was not computed, because the scoring scripts of the paper do not implement it. TM-score is reported for all arms in scores_v3/ (monomer medians 0.992, 0.989 and 0.983).

## References

1. Abramson J, Adler J, Dunger J, Evans R, Green T, Pritzel A, et al. Accurate structure prediction of biomolecular interactions with AlphaFold 3. Nature. 2024;630:493–500. doi:10.1038/s41586-024-07487-w

2. OpenFold Consortium. OpenFold3-preview-2 technical report. 2025. Available from: https://portal.openfold.io/reports/of3p2_technical_report.pdf

3. Wohlwend J, Corso G, Passaro S, Getz N, Reveiz M, Leidal K, et al. Boltz-1: democratizing biomolecular interaction modeling. bioRxiv [Preprint]. 2024. doi:10.1101/2024.11.19.624167

4. ByteDance AML AI4Science Team, Chen X, Zhang Y, Lu C, Ma W, Guan J, et al. Protenix: advancing structure prediction through a comprehensive AlphaFold3 reproduction. bioRxiv [Preprint]. 2025. doi:10.1101/2025.01.08.631967

5. Škrinjar P, Eberhardt J, Tauriello G, Schwede T, Durairaj J. Have protein– ligand co-folding methods moved beyond memorisation? bioRxiv [Preprint]. 2025. doi:10.1101/2025.02.03.636309

6. Dobson L, Tusnády GE, Tompa P. Regularly updated benchmark sets for statistically correct evaluations of AlphaFold applications. Brief Bioinform. 2025;26:bbaf104. doi:10.1093/bib/bbaf104

7. Durairaj J, Adeshina Y, Cao Z, Zhang X, Oleinikovas V, Duignan T, et al. PLINDER: the protein-ligand interactions dataset and evaluation resource. bioRxiv [Preprint]. 2024. doi:10.1101/2024.07.17.603955

8. Neupane P, Liu J, Cheng J. Improving AlphaFold3 by engineering MSA and template inputs. bioRxiv [Preprint]. 2026. doi:10.64898/2026.04.22.720119

9. Xu S, Feng Q, Qiao L, Wu H, Shen T, Cheng Y, et al. Benchmarking all-atom biomolecular structure prediction with FoldBench. Nat Commun. 2026;17:442. doi:10.1038/s41467-025-67127-3

10. Mirdita M, Schütze K, Moriwaki Y, Heo L, Ovchinnikov S, Steinegger M. ColabFold: making protein folding accessible to all. Nat Methods. 2022;19:679–682. doi:10.1038/s41592-022-01488-1

11. ColabFold MSA server database history [Internet]. GitHub; [cited 2026 Nov 18]. Available from: https://github.com/sokrypton/ColabFold/wiki/MSA-Server-Database-History

12. Mirabello C, Wallner B. DockQ v2: improved automatic quality measure for protein multimers, nucleic acids, and small molecules. Bioinformatics. 2024;40:btae586. doi:10.1093/bioinformatics/btae586

13. Buttenschoen M, Morris GM, Deane CM. PoseBusters: AI-based docking methods fail to generate physically valid poses or generalise to novel sequences. Chem Sci. 2024;15:3130–3139. doi:10.1039/D3SC04185A

14. Škrinjar P, Eberhardt J, Studer G, Tauriello G, Schwede T, Durairaj J. Evaluating generalization in protein–ligand cofolding methods. Nat Struct Mol Biol. 2026;33:782–794. doi:10.1038/s41594-026-01797-5

15. [dataset] wwPDB consortium. Protein Data Bank: the single global archive for 3D macromolecular structure data. Nucleic Acids Res. 2019;47:D520–D528. doi:10.1093/nar/gky949

16. [dataset] Soultana M, Odesanya OI, Gaitho M, Richman G. Data and code for: OpenFold3 often receives a benchmark target’s own structure as a template. Zenodo. 2026. doi:10.5281/zenodo.24517157

17. Google DeepMind. AlphaFold [software]. GitHub; [cited 2026 Nov 23]. Available from: https://github.com/google-deepmind/alphafold (setting -- max_template_date)

